# A beneficial bacterium influences *myo*-inositol homeostasis to protect plants during drought

**DOI:** 10.64898/2026.09.03.748652

**Authors:** Tri Tran, Kevin D. Santiago-Morales, Bridget O’Banion, Brittni R. Kelley, Sarah Lebeis

## Abstract

Exposure to abiotic stress is one of the primary factors limiting crop productivity with drought stress is the most prevalent. Under drought, plants can produce osmolytes that increase water retention and prevent severe drought symptoms. Plant responses by extension also impact their associated root microbiomes through altered root metabolite concentrations and exudation. For example, myo-inositol (MI) serves as a precursor to osmolytes and also serves as a mediator for plant-microbe interactions in *Arabidopsis thaliana* (Arabidopsis). Here, we inoculated plants with *Pantoea sp.* R4 (R4), an isolate that can catabolize MI, and subjected them to drought. We observed that R4 colonized plants experiencing drought maintained their leaf relative water content while uninoculated planted did not, and that this colonization coincided with enrichment of MI in the shoots and depletion of MI in the roots. Interestingly, exogenous MI alone rescued water retention in wild-type Col-0, but not in *int1* mutants, which lack the tonoplast MI transporter. Together, our results show that R4 colonization under drought conditions increases MI in the shoots, that this accumulation is associated with increased leaf water retention, and that INT1-mediated MI transport is required for this protection. Our results suggest that microbial colonization can alter how MI is localized in plants, which can inform future development of treatments to protect plants from drought.

## Introduction

Drought is one of the most significant threats to global agriculture, and climate change is expected to make water scarcity more frequent and severe, threatening crop productivity and food security worldwide (Intergovernmental Panel On Climate Change (IPCC), 2023). When plants experience water deficit, they respond physiologically by reducing cell expansion and division, slowing photosynthesis, and adjusting to reduced nutrient availability (Chaves et al., 2003). Drought reduces plant metabolic rates and normal growth through stomatal closure to limit water loss, which restricts carbon assimilation and can promote the buildup of reactive oxygen species (ROS), causing oxidative damage (Chaves et al., 2009). Prolonged drought reduces crop yields and, in severe cases, causes crop failure.

To cope with these physiological effects, plants activate molecular mechanisms to support survival under water-deficient conditions. Central to this response is the phytohormone abscisic acid (ABA), which accumulates under water deficiency and acts as a primary signal coordinating drought responses, such as stomatal closure and altered gene expression (Sah et al., 2016; Takahashi et al., 2020). Among plant genes with increased expression under drought are those responsible for production of common plant osmoprotectants include proline, glycine betaine, and sugars or sugar alcohols such as trehalose and mannitol (Ashraf & Foolad, 2007; Shinozaki & Yamaguchi-Shinozaki, 2006). Among these osmolytes, myo-inositol (MI) also occupies a central position because it serves as a precursor to a range of downstream protective compounds, including galactinol and the raffinose-family oligosaccharides (Loewus & Murthy, 2000), and because MI and its derivatives act as osmoprotectants under stress (Valluru & Van Den Ende, 2011).

In Arabidopsis, overexpression of maize MYO-INOSITOL PHOSPHATE SYNTHASE led to higher leaf MI concentrations which increased drought tolerance (Li et al., 2020). Moreover, the subcellular location of MI can play a role on its availability throughout the plant and within cells. For example, the tonoplast transporter INOSITOL TRANSPORTER 1 (INT1) exports MI from the vacuole to the cytosol, contributing to how MI is distributed and stored within the cell (Schneider et al., 2008). Exogenous MI application alleviates water-deficit stress in mungbean, pepper, and maize, improving leaf water status and limiting oxidative damage (Liu et al., 2025; Patel et al., 2025; Yildizli et al., 2018), indicating that several crop species benefit from increased MI availability. However, if MI availability protects plants directly, as an osmoprotectant precursor, or a signaling molecule remains unclear (Gillaspy, 2011, Perera et al., 2008).

Plants can recruit and select for microbes that offer additional drought protection via the root exudation of metabolites that shape the composition of the rhizosphere microbiome, selectively recruiting beneficial microbes that can support host stress tolerance (Trivedi et al., 2022). Drought alters this exuded pool, increasing the exudation of sugars and sugar alcohols, including MI (Canarini et al., 2016), and shifts the microbial community toward taxa that persist under low water availability. Like plants, these bacteria synthesize their own osmoprotectants, including trehalose, to maintain cellular osmotic balance during drought stress (Vílchez et al., 2016). Bacteria also use strategies unique to microbes, such as forming protective biofilm on surfaces through extracellular polymeric substances (EPS) production (Trivedi et al., 2022). These adaptations support both bacterial survival and the microbial communities surrounding plant roots, improving soil structure and water availability for the plant.

Our lab has previously characterized *Pantoea* sp. R4 (R4), a bacterial isolate with multiple plant-growth promoting traits that colonizes Arabidopsis robustly and produces EPS (Moccia et al., 2020; O’Banion et al., 2023). Notably, MI serves as a carbon source, contributes to biofilm production (Supplemental Figure 3), and potentially a signaling molecule for R4 (O’Banion et al., 2023). Plant MI transport positively impacts R4 endophytic colonization, linking this plant-microbe interaction directly to MI metabolism (O’Banion et al., 2023). This suggests that plant MI pools impact their microbiota but opens the question of whether endophytic microbes influence MI levels in plant tissues. This would be especially relevant to understand in the context of drought, where MI is involved in plant responses to this abiotic stressor.

Here, we investigate how *Pantoea* sp. R4 affects MI accumulation and distribution in *Arabidopsis thaliana* under drought, and whether this interaction shapes the plant’s ability to withstand water deficit. Using R4 inoculation, exogenous MI and ABA treatments, and an Arabidopsis *int1* mutant, we show that microbial colonization leads to MI accumulation in the shoots, and that INT1-mediated MI transport is required for MI-associated drought protection.

## Materials and Methods

*Germination, inoculation, and growing conditions of Col-0 with* Pantoea *sp. R4 Arabidopsis thaliana* ecotype Col-0 seeds were surface sterilized, stratified, and germinated for 7 days on ½-strength Murashige and Skoog (MS) basal medium lacking *myo*-inositol (MP Biomedicals, cat. no. 2623022) supplemented with 0.8% (w/v) Phytoagar (PlantMedia, cat. no. 40100072) and 1% (w/v) sucrose. Plates were maintained in a growth chamber at 24 °C/22 °C (day/night) under long-day conditions (16 h light/8 h dark) with a photosynthetic photon flux density of approximately 120 µmol m ² s ¹. On day 8, seedlings were transferred to ¼-strength MS basal medium without myo-inositol solidified with 0.6% (w/v) Phytoagar, whose surface had been pretreated either with 150 µL sterile 1× phosphate-buffered saline (PBS; prepared from 10× PBS, Fisher Scientific, cat. BP399-20) as a no-bacteria control (NB) or with 150 µL of a 10 CFU/mL of *Pantoea* sp. R4 in 1× PBS. Seedlings were grown for a further two weeks under the same growth conditions before being transferred to pots.

After two weeks, three subsets of seedlings were harvested to quantify R4 colonization at this stage by determining colony-forming units (CFUs). Seedlings from each subset were weighed, rinsed with sterile 1× PBS, homogenized, and the resulting homogenates were serially diluted and plated on MacConkey agar. The mean bacterial load across the three samples was 3.19 × 10^6 CFU g⁻¹ of fresh plant tissue.

The remaining 24 healthy seedlings from the germination medium were transplanted into sterile pots containing 100 g of double-autoclaved growth medium composed of 4:1 ratio of Sure-Mix (Michigan Grower Products, Inc.) potting soil and sand mixture and grown for an additional two weeks in a growth chamber under the same conditions described above. All plants were watered daily with 10 mL sterile water for 14 days during this establishment period. After establishment, six plants per treatment (NB and R4-inoculated) were assigned either to continued daily watering with 10 mL sterile water or to drought, imposed by complete withholding of water, yielding four treatment groups: NB-watered, NB-drought, R4-watered, and R4-drought. Plants were harvested for fresh root and shoot tissues 10 days after initiation of the drought treatment, and pot weights in watered and drought treatments were monitored throughout to assess water loss rates across treatments.

### Plant harvests and Pantoea sp. R4 CFU determinations

Plants from both watered and drought treatments were harvested at the onset of drought symptoms in the drought treatment group. At harvest, final pot weights were obtained and a single leaf from each plant was collected to determine relative water content (RWC). Bolting shoots were trimmed and removed, leaving only rosettes for harvesting. Using a flame-sterilized glass pan and tweezers, plants were gently removed from pots and soil was separated from the roots. Flame-sterilized scissors were used to separate rosettes and roots, with rosettes placed in 5 ml tubes and snap frozen with liquid nitrogen to use for MI measurements. Roots were placed in 50 mL tubes with 25 mL of sterile harvest phosphate buffer + 0.01% Silwet (HPB) (Lundberg et al., 2012) and vortexed for 20 seconds. Flame sterilized forceps were used to transfer roots to sterile 5 mL tubes with 4 mL phosphate buffer and placed in a bath sonicator for 10 minutes. Following bath sonication, roots were gently removed from the tubes, soil debris was removed from the tube, and roots were returned to the same tube. Roots were washed twice more with sterile phosphate buffer in the same manner. Following the final wash, excess water was shaken off the roots and these were placed in a new sterile 5 ml tube to obtain fresh weight before storing at 4°C overnight. Roots were then homogenized with garnet beads and 1 mL sterile 1X PBS in a Tissue Lyser 2 (QIAGEN cat. no. 85300) at 30 s^-1^ for 10 minutes. Samples were serially diluted to 10^-4^ and spread on MacConkey plates to select for Gram-negative bacteria and incubated at 28°C for 24 hours. Plates were removed from the incubator and left to incubate at room temperature for 48 hours before enumerating.

CFUs were also determined from no plant control samples by placing 5 mL soil volume from individual pots into 15 mL tubes. Seven mL of phosphate buffer was added to each tube before vortexing for 20 seconds and allowing the soil to settle. The resulting supernatant (approximately 4 mL) was transferred to a labeled sterile 5 mL tube, centrifuged at 4,000 rpm for 10 minutes, and placed at 4°C overnight before serial dilution and plating with root tissue samples as described above.

### Relative water content (RWC) measurements

To determine RWC, single leaves collected from each plant were weighed immediately after collection to obtain fresh weight (FW) using an analytical balance. Leaves were then floated in sterile distilled water in covered petri dishes and rehydrated in the dark at 4°C for approximately 24 hours to obtain turgid weight (TW) by blotting the leaf surface dry before weighing. Leaves were then oven-dried at 65°C for approximately 48 hours to obtain dry weight (DW). RWC was then calculated using (FW-DW)/(TW-DW) × 100.

### Extraction and measurement of myo-inositol concentrations

*Myo*-inositol (MI) was extracted separately from roots and shoots with methanol containing *myo*-inositol-C-d6 as an internal standard (MilliporeSigma, cat. 616184), and samples were analyzed by ultra-performance liquid chromatography–tandem mass spectrometry (UPLC–MS/MS) on a Xevo TQ-XS triple quadrupole mass spectrometer (Waters Corporation, Milford, MA) at the MSU Metabolomics Core Facility. Briefly, 100 mg of lyophilized tissue was extracted in ice-cold buffer consisting of 80:20 (v/v) methanol:water, 0.1% formic acid, 0.1 g/L butylated hydroxytoluene (BHT), and 10 nM internal standard. Suspensions were incubated at 4 °C with gentle shaking for 16 h, then filtered and stored at −80 °C until analysis. Data was processed using MassLynx software (TargetLynx XS module, Waters).

### RNA extraction, library construction and sequencing

Total RNA was extracted from frozen root tissue using the RNeasy PowerSoil Total RNA Kit (Qiagen, cat. 12866). Three biological replicates per treatment were processed. Frozen tissues were carefully homogenized, and approximately 3 g of material was transferred to a bead tube provided with the kit, followed by RNA isolation according to the manufacturer’s instructions. The resulting RNA pellet was resuspended in 25 µL RNase/DNase-free water. To remove residual genomic DNA, RNA samples were treated with the RNase-Free DNase Set (Qiagen, cat. 79254) according to the manufacturer’s protocol. Nucleic acid concentrations were determined using a Qubit 2.0 Fluorometer (Life Technologies). Samples were stored at −80 °C prior to submission to the U.S. Department of Energy Joint Genome Institute (JGI) for sequencing.

For RNA sequencing, stranded cDNA libraries were prepared using the Illumina TruSeq Stranded RNA LT kit (Illumina). Ten nanograms of total RNA were fragmented using divalent cations at elevated temperature, followed by first-strand cDNA synthesis with random hexamer primers and SuperScript II reverse transcriptase (Invitrogen).

Second-strand synthesis was then performed to generate double-stranded cDNA. The resulting cDNA was subjected to end repair, A-tailing, adapter ligation, and eight cycles of PCR amplification. Libraries were quantified using KAPA Library Quantification Kits (KAPA Biosystems) and a Roche LightCycler 480 real-time PCR instrument. Sequencing was carried out on an Illumina NovaSeq platform using NovaSeq XP v1 reagent kits and an appropriate flow cell configuration, following a sample-dependent indexed run recipe

### RNA-seq analysis

Quality-controlled and filtered RNA sequencing data was downloaded from the Joint Genome Institute’s Genome Portal. Transcript abundances were determined using kallisto (Bray et al., 2016). A reference index was built using the Arabidopsis Thaliana Reference Transcript Dataset 2, updated for use in Quantification of Alternatively Spliced Isoforms (AtRTD2-QUASI) (Zhang et al., 2017). Transcript quantification was performed using kallisto quant with 100 bootstraps and the default seed of 42. The outputs were then imported into sleuth for differential expression analysis at the transcript isoform level (Pimentel et al., 2017). A likelihood-ratio test was used to determine significance values, and a wald test was used to determine b-values (analogous to log2FC). We performed a PANTHER overrepresentation test (Fisher’s exact test with Bonferroni correction) on the output gene lists using the PANTHER (v19.0) classification system and GO Ontology Database (released 2025-07-22).

### Biofilm Production Assay

*Pantoea* sp. R4 was inoculated into Luria Bertani (LB) broth cultures and incubated with shaking until turbid (12-18 hours) at 28°C. Optical density at 600nm (OD600) for each culture was then measured. Cultures were diluted to a final OD600 of 0.05. Diluted cultures were washed three times in sterile 1X M9 minimal salts media supplemented with 10mM either glucose or *myo*-inositol, or 5mM each glucose and *myo*-inositol. 100µL of each culture as well as media blanks were transferred to wells of a Corning Falcon 96-well flat bottom clear plate. The plate was incubated at 28°C without shaking for 48 hours. The OD600 was collected. The plate was gently shaken out to remove planktonic bacteria and rinsed by submersion in DI water. The rinse was gently shaken out until no liquid remained in the wells. 125 µL of 0.1% crystal violet were added to each well and allowed to stain for 10 minutes. The plate was gently shaken out and rinsed by submersion in DI water again, then shaken out until no liquid remained in the wells. The plate was left upright to dry overnight. 200uL of 30% acetic acid were added to each well and allowed to solubilize the crystal violet for 10 minutes before mixing by pipetting up and down. The OD595 of the resuspended solution was collected. OD595 values for appropriate blanks were subtracted from sample values before reporting.

### Germination, growth conditions, and chemical treatments of Col-0 and int1 plants

Arabidopsis thaliana genotypes Col-0 and int1 were surface sterilized, stratified, and germinated for two weeks on sterile vertical plates without microbial inoculation. After two weeks, 30 healthy seedlings per genotype were selected and transferred to double-autoclaved growth medium consisting of a 4:1 Sure-Mix:sand mixture. All plants were well-watered with 10 mL sterile water daily for two weeks. Established plants were then divided into three treatment groups (10 plants per group) receiving either a sterile water treatment, 10 mM myo-inositol (Thermo Scientific Chemicals, cat. J60828.22), or 100 µM abscisic acid (ABA; Thermo Scientific Chemicals, CAS 21293-29-8). Drought was imposed on a subset of five seedlings per treatment group by complete water withholding beginning two days after chemical application, while the remaining seedlings continued to receive 10 mL sterile water daily. All plants were harvested 10 days after initiation of the drought treatment, and roots and shoots were collected for downstream analyses.

### Statistical analyses

All analyses for physiological data were done in GraphPad PRISM (10.6.1). Normality was tested for all datasets, and two-way ANOVAS were used to determine differences across treatments. Tukey test multiple comparisons were done post-hoc ANOVA for all datasets and treatments, and differences with p<0.05 were reported. For the genotype × water treatment × chemical treatment experiment, a three-way ANOVA was done to determine the significance of each treatment (supplemental fig. 3). Post-hoc multiple comparisons of the three-way ANOVA were also done using a Tukey test. *Pantoea* sp. R4 colonization data was analyzed with a Welch’s unpaired t-test.

## Results

### Pantoea sp. R4 confers Arabidopsis resistance to drought

To determine whether *Pantoea* sp. R4 inoculation impacts drought tolerance and plant physiology, relative water content (RWC), dry biomass, and colonization rates of *Pantoea* sp. R4 were measured in Col-0 plants following water withholding (Supplemental Figure 1). Additionally, *Pantoea* sp. R4 colonization was confirmed after two weeks of colonization on plates prior to transplanting seedlings into pots with soil.

After 10-14 days of water-withholding, leaves were visibly wilting on uninoculated plants while R4-inoculated plants showed no phenotypic signs of drought stress (Figure 1A). As expected, drought treatment significantly reduced RWC in uninoculated plants compared to their watered controls (p<0.0001). Contrastingly, RWC remained similar in R4-inoculated plants under both watered and drought conditions. Although there was no significant difference in RWC between inoculated and uninoculated plants under well-watered conditions, R4 inoculated plants exhibited significantly higher RWC compared to uninoculated plants under drought conditions (p<0.001) (Figure 1B). Confirming that the effect of drought on RWC was dependent on inoculation, two-way ANOVA showed significant main effects of water treatment (p<0.0001) and inoculum (p<0.01), as well as a significant interaction between water treatment and inoculum (p<0.001). Together, these results suggest that R4 inoculation confers drought tolerance in plants by maintaining leaf RWC under water deficit.

**Figure 1.**
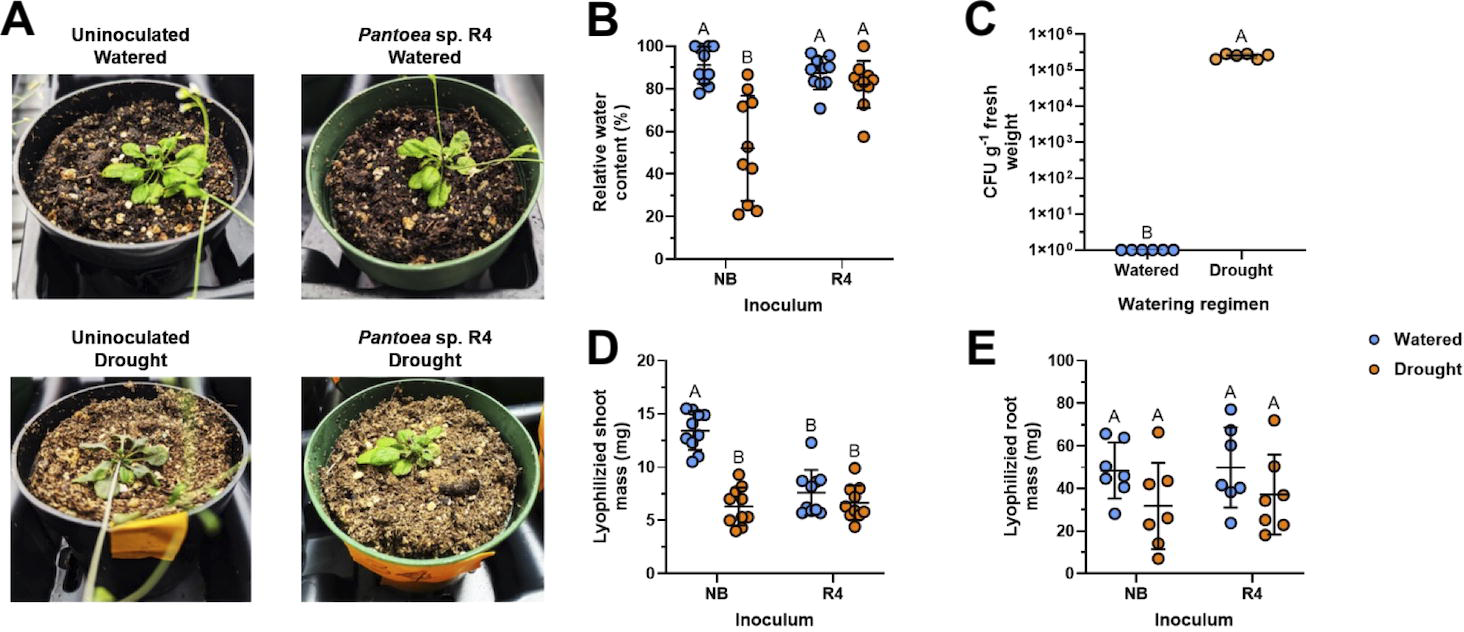
Arabidopsis inoculated with *Pantoea* sp. R4 maintain leaf relative water content under drought stress. (A) Representative plants from each treatment group at the end of the experiment, with well-watered plants on top and drought plants on the bottom, and uninoculated plants on the left and inoculated plants on the right. (B) RWC of leaves in Col-0 plants subjected to well-watered and drought conditions with or without *Pantoea* sp. R4 inoculation. (C) Recovery of *Pantoea* sp. R4 in inoculated plants under well-watered and drought conditions expressed as colony forming units per gram of fresh root weight (CFU g⁻¹ FW) at the end of the experiment. (D) Dry shoot mass and (E) root mass of Col-0 plants across inoculum and water treatments. For panels B-E, data are presented as individual points with mean and standard deviation. For panels B and D, n=10, for panel E, n=7, and for panel C, n=6. Statistical significance for panels B, D, and E was determined by two-way ANOVA followed by Tukey’s multiple comparisons test. Panel C was analyzed using an unpaired t-test with Welch’s correction. Different letters indicate statistically significant differences between groups (p<0.05).

To determine if R4 colonization was affected by drought, we used MacConkey selective media to quantify its root colonization levels at after 2 weeks (Supplemental Figure 2) and the end of our experiment (Figure 1C). Under well-watered conditions, no colonies of R4 were recovered at the end of the experiment, indicating that R4 was unable to persist endophytically. Notably, colonization levels were lower at the end of the experiment in soil (Figure 1C) than they were after 2 weeks on plates (Supplemental Figure 2). In contrast, R4 robustly colonized plants under drought conditions (p<0.0001) (Figure 1C). These results demonstrate that continued R4 colonization in this context is drought-dependent.

To assess if drought treatment or *Pantoea* sp. R4 inoculation affected plant biomass, mass of lyophilized shoot and root tissues were measured. Dry shoot mass was significantly higher in the uninoculated well-watered plants than in any other group (p<0.0001) (Figure 1D). This indicates that both drought and R4 colonization independently reduced shoot biomass, with a significant interaction between the two factors (p<0.0001). In contrast, root mass showed no significant differences across any group (Figure 1E). Together, these results indicate that imposed drought negatively impacted uninoculated plants while plants inoculated with *Pantoea* sp. R4 maintained RWC in their leaves.

### Drought resistance provided by Pantoea sp. R4 correlates with altered myo-inositol levels in root and shoot tissues

To determine if our experimental drought treatment impacted MI homeostasis in roots, we performed RNA sequencing on uninoculated root tissue and examined expression levels for genes known to be associated with MI cycling in plant cells (Figure 2A). We observed that the majority of genes in MI cycling demonstrated higher expression levels in well-watered plants than droughted plants (Figure 2A). To establish how these results translated into MI level in plants under our drought conditions, we measured MI concentration in roots, as well as shoots. Root MI concentrations showed a significant 7.6-fold increase under drought in uninoculated plants compared to watered controls (p<0.05) (Figure 2B), suggesting that the lower expression of MI cycling-related genes resulted in higher accumulation of MI in roots. Interestingly, there was no significant difference in leaf MI concentration between uninoculated well-watered and drought treatments (Figure 2C).

**Figure 2:**
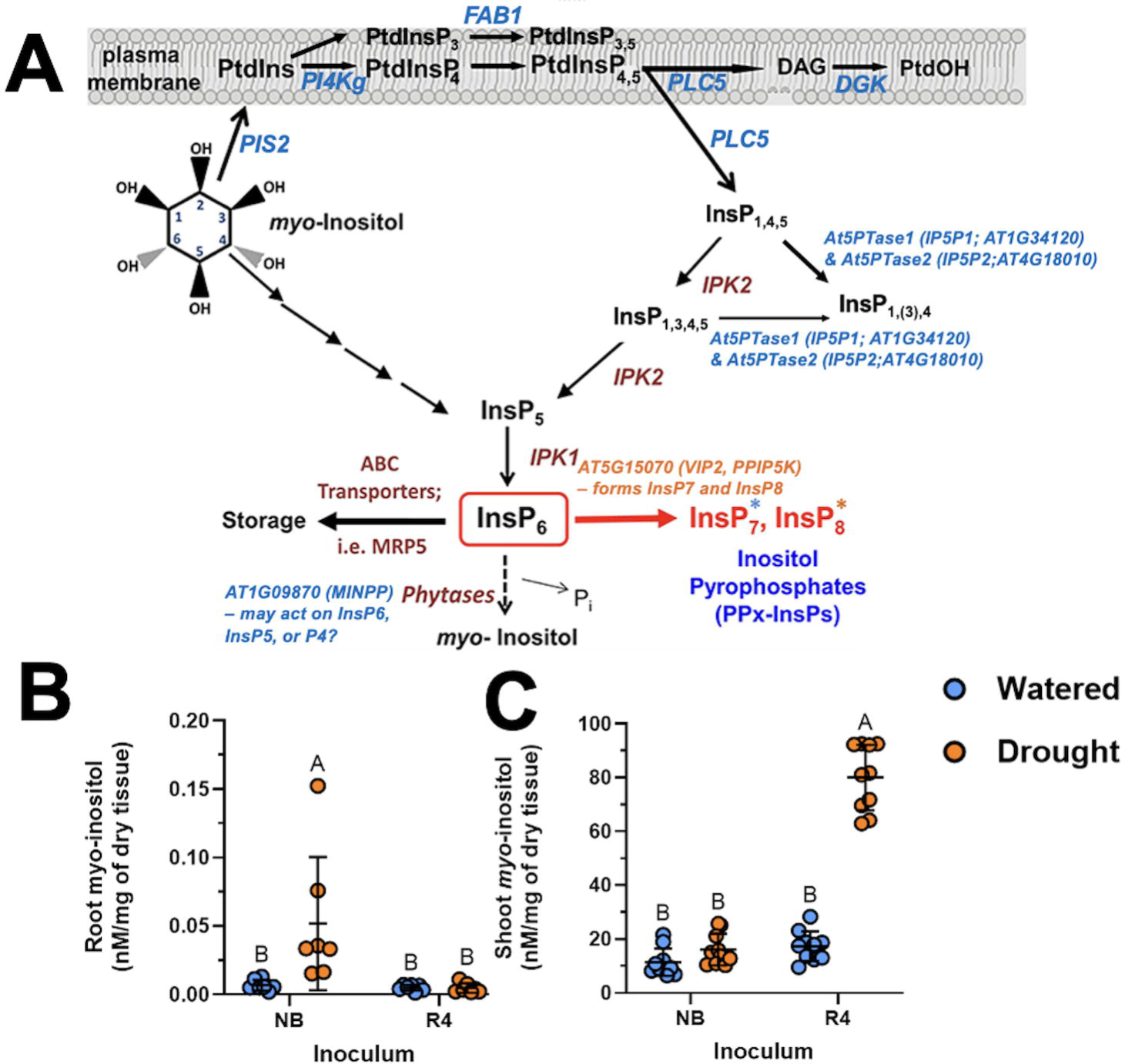
***Pantoea* sp. R4 inoculation alters Col-0 *myo*-inositol levels in shoots and roots in drought conditions.** (A) Root expression of genes involved in *myo*- inositol (MI). Genes with decreased expression under drought conditions (p-val <0.05) are highlighted in blue. Genes with increased expression under drought conditions (p- val <0.05) are highlighted in orange. (B) Root MI and (C) Shoot MI concentrations in Col-0 plants inoculated with R4 or left uninoculated (NB) under well-watered or drought conditions. Data are presented as individual points with mean and standard deviation. n=7 for (A) and n=10 for (B). Statistical analysis was performed by two-way ANOVA followed by Tukey’s multiple comparisons test. Different letters indicate statistically significant differences between groups (p<0.05).

Because R4 colonization levels are positively impacted by MI transport (O’Banion et al., 2023), we reasoned that R4-mediated drought tolerance might be linked to changes in MI distribution. We measured shoot and root MI concentrations to determine if R4 colonization also affected MI distribution across plant tissues and if this could correlate to R4 plant colonization under drought conditions. While we observed no difference in MI levels between watered and drought-stressed R4-inoculated root samples (Figure 2B), R4-inoculated plants showed a significant 4.6-fold increase in shoot MI under drought (p<0.0001) (Figure 2C). A significant interaction between watering treatment and inoculum was also observed for root MI (p<0.05). Together, these results show that R4 shifts localization of drought-induced MI accumulation as uninoculated plants build up MI in roots under drought, while R4-inoculated plants instead accumulate MI in shoots. This redistribution toward the shoot coincides with the maintained leaf RWC seen in R4-inoculated plants, suggesting that R4 supports drought tolerance by promoting MI accumulation in above-ground tissues where it can help retain water.

### INT1-dependent myo-inositol drought protection

To dissect the role of MI and its transport in Arabidopsis responses to drought stress, we exogenously added 10 mM MI to uninoculated watered and drought plants. We were also interested in determining if inositol vacuolar transport affected MI-driven responses to drought, so we included *int*1 lines in these water treatments. Furthermore, we exposed Col-0 and *int*1 plants to 100 µM exogenous ABA to determine if the MI-derived response to drought was related to a general stress response driven by this phytohormone.

Overall, the RWC in Col-0 plants revealed a significant interaction between chemical treatments and watering regimes (p<0.0001). In the leaves of Col-0 experiencing drought, we observed the expected decrease in RWC with no chemical treatment (Figure 3A). Notably, Col-0 plants treated with MI on their roots maintained equivalent leaf RWC under both watered and drought conditions, indicating that exogenous MI fully protects against drought-induced water loss in Col-0 (Figure 3A). As expected, exogenous ABA root application conferred stable RWC under drought compared to the control treatment, was not as effective as MI (p<0.0001) (Figure 3A), suggesting that exogenous MI To determine if MI transport is required for this protection, we applied chemical treatments to *int1* plants prior to onset of drought. We again observed decrease in RWC in control treatments, which was ameliorated with ABA treatment (Figure 3B). Contrary to what was observed in Col-0, exogenous MI did not rescue or maintain RWC, as a significant reduction was observed between watered and drought conditions (p<0.0001) (Figure 3B). ABA also did not rescue RWC under drought conditions. This confirms that vacuolar transport of MI via INT1 is required for MI-mediated protection against drought.

**Figure 3.**
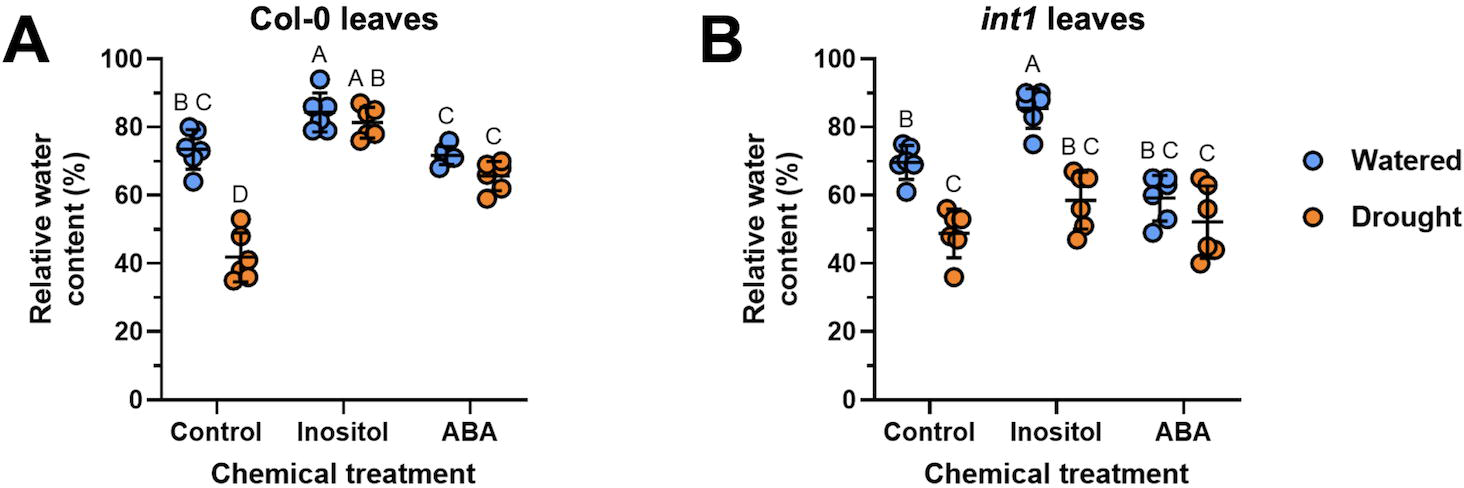
Exogenous *myo*-inositol rescues drought-induced water loss in Col-0 but not in *int1* plants. RWC of leaves from (A) Col-0 and (B) *int1* plants treated with control, 10 mM MI, or 100 µM ABA under well-watered and drought conditions. Data are presented as individual points with mean and standard deviation (n=6). Statistical significance was determined separately for each genotype by two-way ANOVA followed by Tukey’s multiple comparisons test across all treatment and watering condition combinations. Different letters indicate statistically significant differences between groups (p<0.05).

To assess the impact of drought and exogenous MI and ABA on plant biomass, lyophilized shoot and root mass were measured in Col-0 and *int1* plants across chemical and watering treatments. Two-way ANOVA of Col-0 shoot biomass revealed significant main effects of chemical treatment (p<0.0001) and water treatment (p<0.001), and a significant interaction between the two (p<0.001). Control watered Col- 0 had significantly higher biomass than all other groups (p<0.0001 for all comparisons), indicating that both drought and chemical treatment independently suppress shoot growth in Col-0 (Figure 4A). Both MI-treated and ABA-treated plants had reduced shoot biomass compared to control drought plants (p<0.01 for MI watered, p<0.0001 for all other chemical groups), indicating that chemical treatment suppressed shoot growth more severely than drought alone.

**Figure 4:**
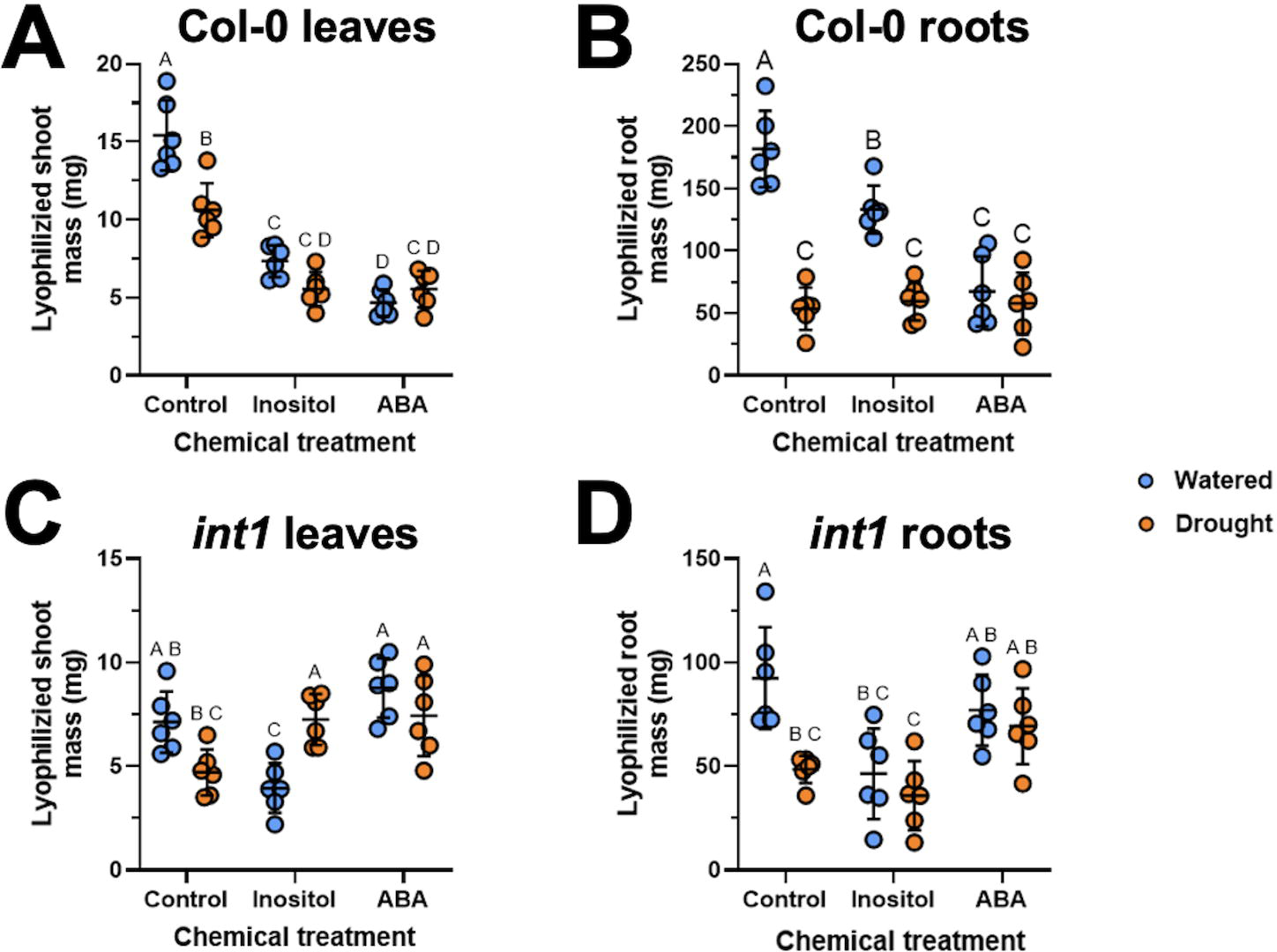
**INT1 mediates biomass under drought stress and chemical treatment**. Lyophilized (A) shoot and (B) root biomass of Col-0 and *int1* (C) shoots and (D) roots across water and chemical treatments. For all panels, data are presented as individual points with mean and standard deviation (n=6). Statistical significance was determined separately for each genotype by two-way ANOVA followed by Tukey’s multiple comparisons test. Different letters indicate statistically significant differences between groups (p<0.05), determined for each genotype.

Col-0 root biomass showed significant main effects of chemical treatment (p<0.0001), water treatment (p<0.0001), and their interaction (p<0.0001). Significant decreases in root biomass were observed between watered and drought plants in the control (p<0.0001) and MI (p<0.0001) treatments, whereas no difference was observed in ABA (Figure 4B). Under watered conditions, both MI and ABA decreased root biomass relative to control (p<0.05 and p<0.0001), whereas under drought no differences were observed among treatments.

In contrast, two-way ANOVA of *int1* shoot biomass revealed a significant interaction between chemical treatment and watering regime (p<0.0001), a significant effect of chemical treatment (p<0.001), but no significant main effect of water treatment. Interestingly, shoots of watered MI-treated *int1* plants had the strongest decrease in biomass (p<0.01) compared to shoots of control, while ABA-treated plants maintained biomass relative to the control (Figure 4C). Droughted int1 plants increased shoot biomass in both the MI and ABA treatments (p<0.05 for both), and biomass was increased in the MI treatment under drought conditions compared to watered MI-treated plants (p<0.01).

Two-way ANOVA of int1 root biomass revealed significant main effects for chemical treatment (p<0.001) and water treatment (p<0.01), with a significant interaction between them (p<0.05). In watered chemical treatments, MI significantly decreased biomass relative to untreated plants (p<0.01), whereas ABA maintained it (Figure 4D). Under drought conditions, ABA-treated *int1* plants maintained root biomass comparable to control, whereas MI-treated int1 plants had significantly lower biomass than ABA-treated plants (p<0.05). Together, these results indicate that biomass responses to MI and ABA are genotype-dependent, and that the disrupted MI distribution observed in *int1* under drought is accompanied by altered biomass allocation patterns compared to Col-0. Furthermore, our *int1* biomass results indicate that the responses mounted by ABA in the plant could be independent of MI transport.

To understand how RWC and biomass measurements related to MI accumulation, we measured shoot and root MI levels in each genotype. In Col-0 shoots, MI levels significantly increased 13-fold between watered and drought MI-treated plants(p<0.0001) but remained consistent across water regimes in the control and ABA treatments (Figure 5A). In Col-0 roots, drought stress increased MI across all drought treatments compared to their corresponding watered treatments with the control treatment showing a 5.7-fold increase (p<0.0001), 4.9-fold increase in the MI treatment (p<0.0001), and 2.2-fold increase in ABA treatment (p<0.01) (Figure 5B).

**Figure 5.**
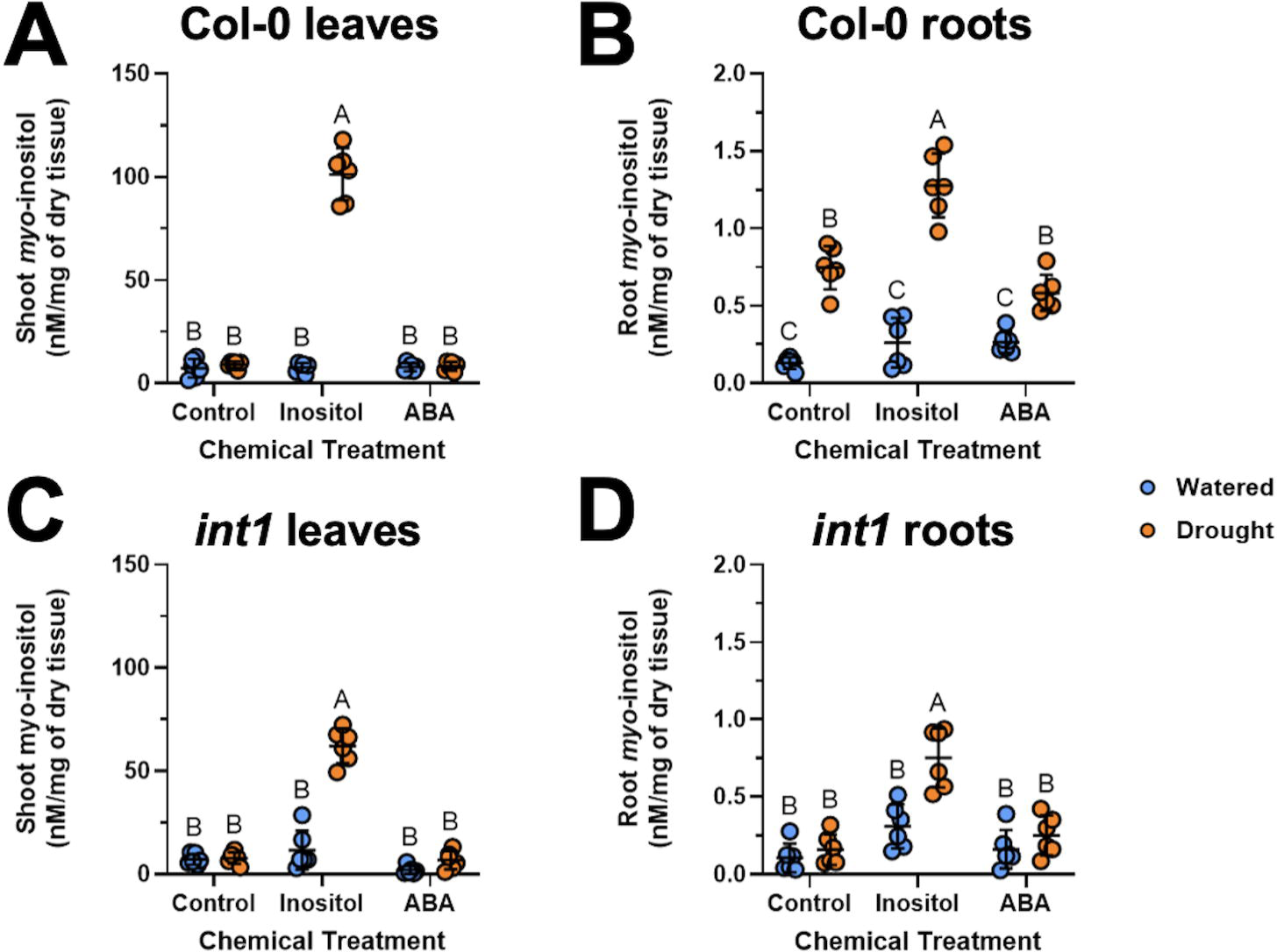
Vacuolar transport of *myo*-inositol via INT1 is required for *myo*-inositol redistribution under drought. (A) Shoot and (B) root MI measurements in Col-0 plants, and (C) shoot and (D) root MI measurements in *int1* plants, under well-watered and drought conditions with control treatments, 10 mM MI, or 100 µM ABA. Data are presented as individual points with mean and standard deviation (n=6). Statistical significance was determined separately for each genotype by two-way ANOVA followed by Tukey’s multiple comparisons test across all treatment and watering condition combinations. Different letters indicate statistically significant differences between groups (p<0.05).

In *int*1, the failure to rescue RWC was reflected in shoot MI levels, which were only elevated 5.3-fold in drought-stressed plants treated with MI (p<0.0001) compared the watered MI treatment (Figure 5C). Unlike Col-0 where drought increased root MI in all treatments, MI levels increased 2.4-fold only in treatments where it was added exogenously to *int1* plants (Figure 5D). Notably, ABA treatment also failed to elevate root MI levels under drought in *int*1, in contrast to Col-0 where ABA significantly increased root MI in drought compared to watered conditions (p<0.01). This suggests that INT1 is needed for MI distribution to roots under drought conditions. Together, these results suggest that MI protects against drought-induced water loss, and that this protection is dependent on INT1-mediated vacuolar MI transport.

## Discussion

Previous work from our lab described that *Pantoea* sp. R4 possesses a wide range of plant growth-promoting traits (Moccia et al., 2020), and that host control of MI mediates R4 colonization and plant-relevant traits, such as motility and EPS production (O’Banion et al., 2023). Here, we extend this characterization to abiotic stress and show that R4 colonization maintains leaf RWC and suppresses the wilting observed in uninoculated plants under drought. Our results suggest that R4 specifically mediates this protection under drought conditions, as its endophytic colonization was only maintained under drought conditions (Figure 1C).

This R4-conferred resilience towards drought was accompanied by a shift in MI accumulation across tissues. Under drought, uninoculated roots decreased expression of genes involved in MI transformation into phosphatidylinositol and phytic acid (Figure 2A), which was accompanied by accumulated MI in roots (Figure 2B). Interestingly, only R4-inoculated plants accumulated it in shoots (Figure 2C), which coincided with leaf water retention (Figure 1B). MI is a recognized osmolyte and its exogenous application partially improves drought tolerance across diverse plants (Liu et al., 2025; Patel et al., 2025; Yildizli et al., 2018). Hence, our observations are consistent with shoot MI accumulation playing a role in above-ground water retention. Specifically, we interpret that redistribution of MI toward the shoot is associated with the R4-mediated protective phenotype. However, whole above- and below-ground tissue MI measurements cannot resolve if this redistribution reflects altered transport, synthesis, or partitioning between subcellular pools, which could be addressed with measurements from subcellular fractions.

Our results from the chemical treatment experiments, along with the *int1* mutant, which lacks the tonoplast-localized transporter that exports vacuolar MI to the cytosol, implicates MI homeostasis as a mechanism involved in drought tolerance in R4- inoculated plants. Previous results from our lab showed that R4 colonized the *int1* mutant (O’Banion et al., 2023) at significantly lower levels than wild-type Col-0, and that this deficit was rescued by exogenous MI application. In our experiments, exogenous MI rescued RWC in Col-0 but not in *int1*, suggesting that the MI transport that promotes R4 colonization may also contribute to the protective effect on water retention. Furthermore, that INT1 is implicated in both phenotypes points toward host cytosolic MI availability, rather than overall MI abundance, as the driver of R4 colonization and drought protection. Despite this, the specific molecular mechanisms linking R4, host MI localization, and water retention remain uncharacterized and present opportunities for future work.

A future step towards this direction would be to determine whether R4 catabolism of MI is associated with its drought protection phenotypes. In the same work by O’Banion et al. (2023), deletion of *iol*G, which encodes the first step of R4 MI catabolism, showed that *iolG* was not essential for root colonization after 1 or 2 weeks (O’Banion et al. 2023 and Supplemental Figure 2). Notably, increased motility in the presence of MI was observed in both wild-type and the Δ*iol*G mutant, indicating that some effects of MI on R4 are independent of its catabolism and may instead reflect bacterial sensing or transport. Determining whether R4 MI catabolism, transport, or sensing becomes relevant under drought, when host MI distribution shifts and R4 colonization become drought-dependent, could expand the results presented here.

Specifically, this would establish whether R4 must metabolize MI to confer the drought protection. Testing transport and sensing separately would then clarify which aspect of the MI interaction is required. Moreover, expanding our osmolyte measurements to include galactinol and raffinose would provide more insight into how MI levels impact the R4-conferred drought resistance. Since these are synthesized from MI, these measurements would determine if MI alone or downstream osmolyte synthesis are responsible for drought protection. Evidence that bacterial catabolism of MI contributes to pathogenic plant-microbe interactions (Hamilton et al., 2021), and that beneficial strains can alter the fate of host MI under drought (Papadopoulou et al., 2026), motivates these future approaches.

Here we show that Pantoea sp. R4 colonizes Arabidopsis roots under drought but not under well-watered conditions, and that this colonization coincides with reduced drought symptoms and maintained RWC. This protection from drought was accompanied by shifts in MI accumulation, as uninoculated plants had higher leaf MI levels, which was not observed in uninoculated roots. Exogenous MI rescued RWC under drought in Col-0 but not in *int1*, indicating that INT1-mediated vacuolar MI transport is required for MI-associated drought protection. Together, these results suggest that host MI allocation is affected by both R4 colonization and drought, and that bacterial colonization can alter osmolyte distribution in plants by an unknown mechanism. Defining this relationship further could inform the use of microbial inoculants to limit drought damage in crops.

