## Supplemental Figures 1-3 for "A beneficial bacterium influences *myo*-inositol homeostasis to protect plants during drought"

### Slide 1
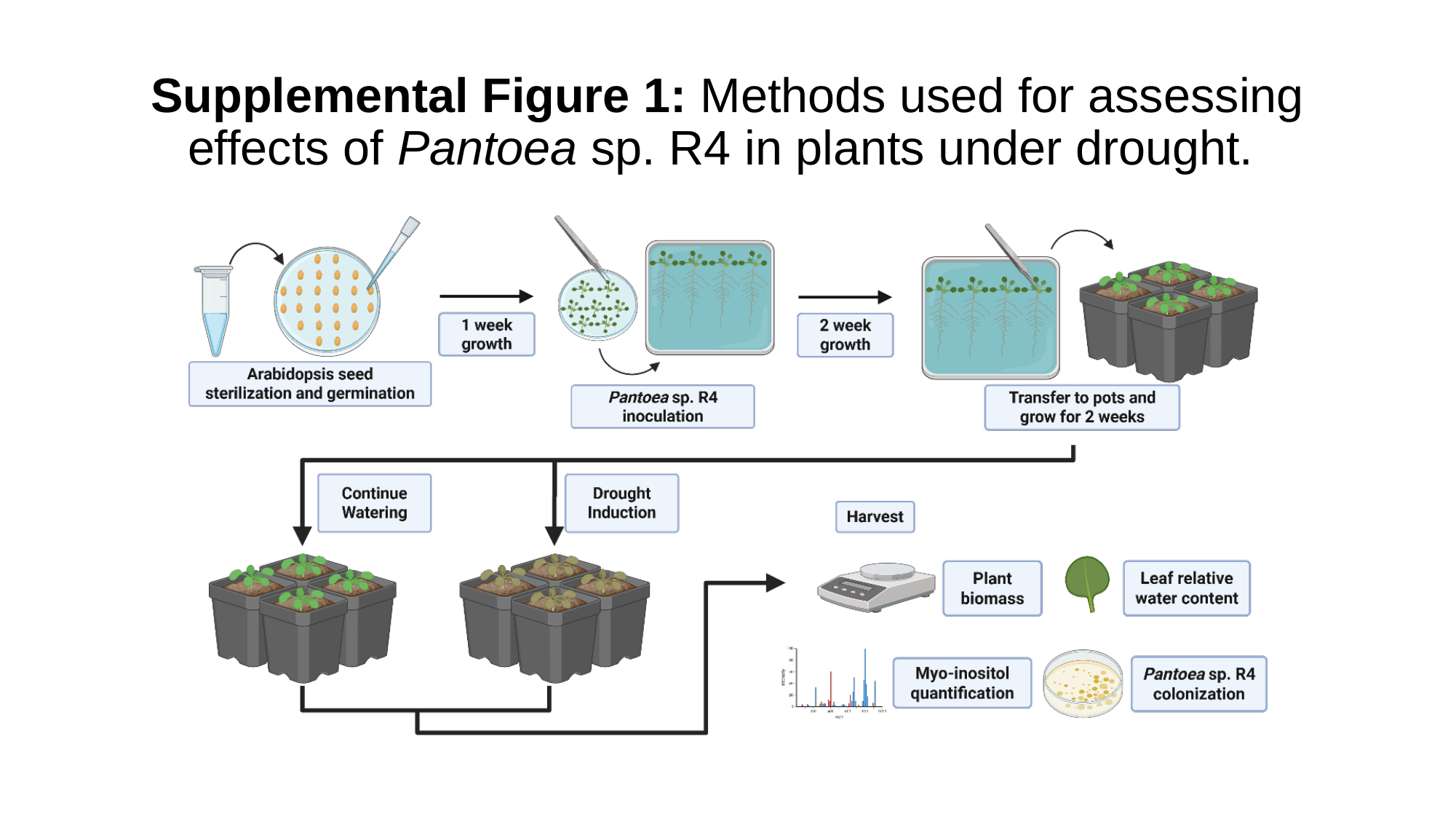

#### Supplemental Figure 1: Methods used for assessing effects of Pantoea sp. R4 in plants under drought.

### Slide 2
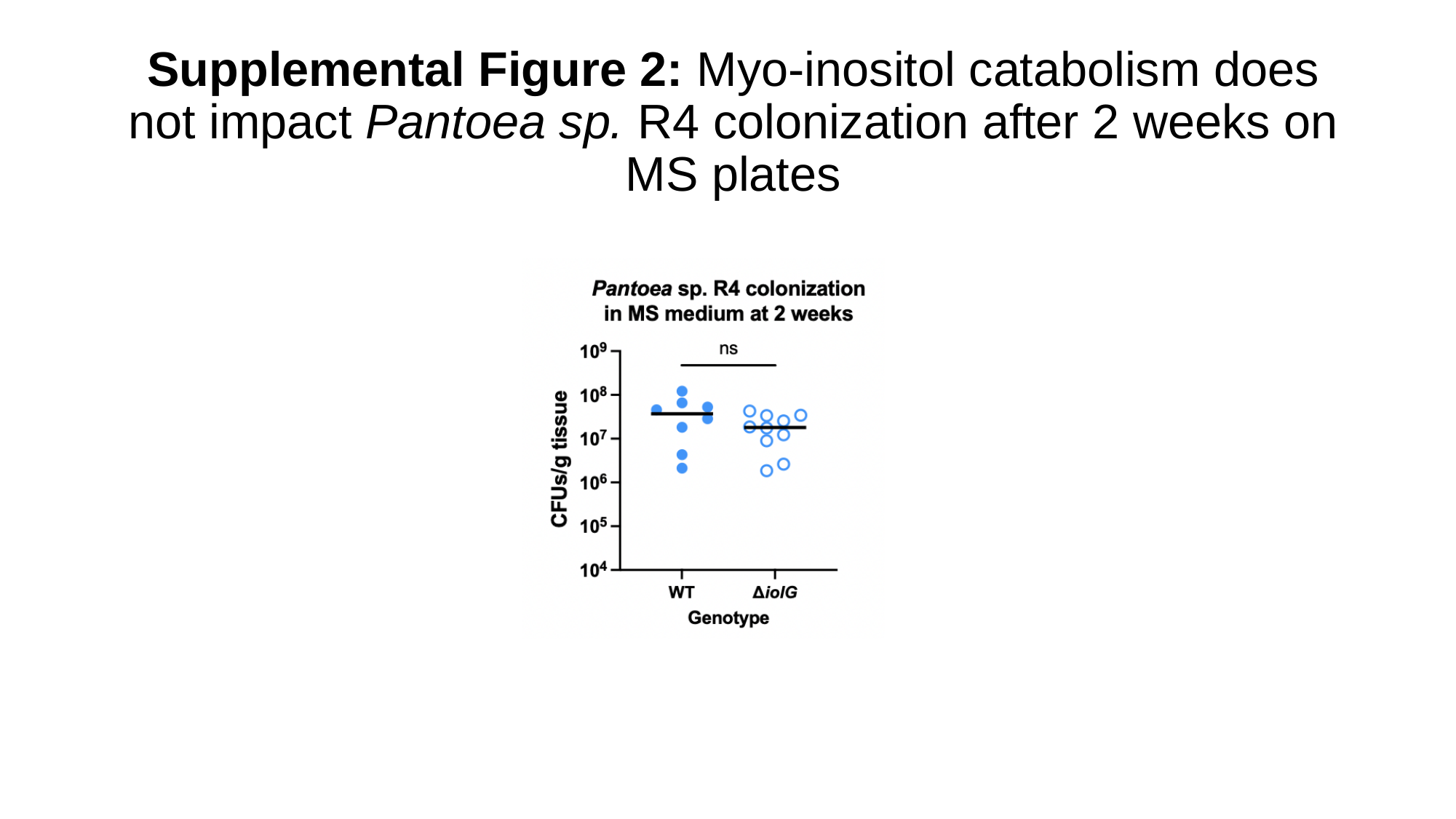

#### Supplemental Figure 2: Myo-inositol catabolism does not impact Pantoea sp. R4 colonization after 2 weeks on MS plates

### Slide 3
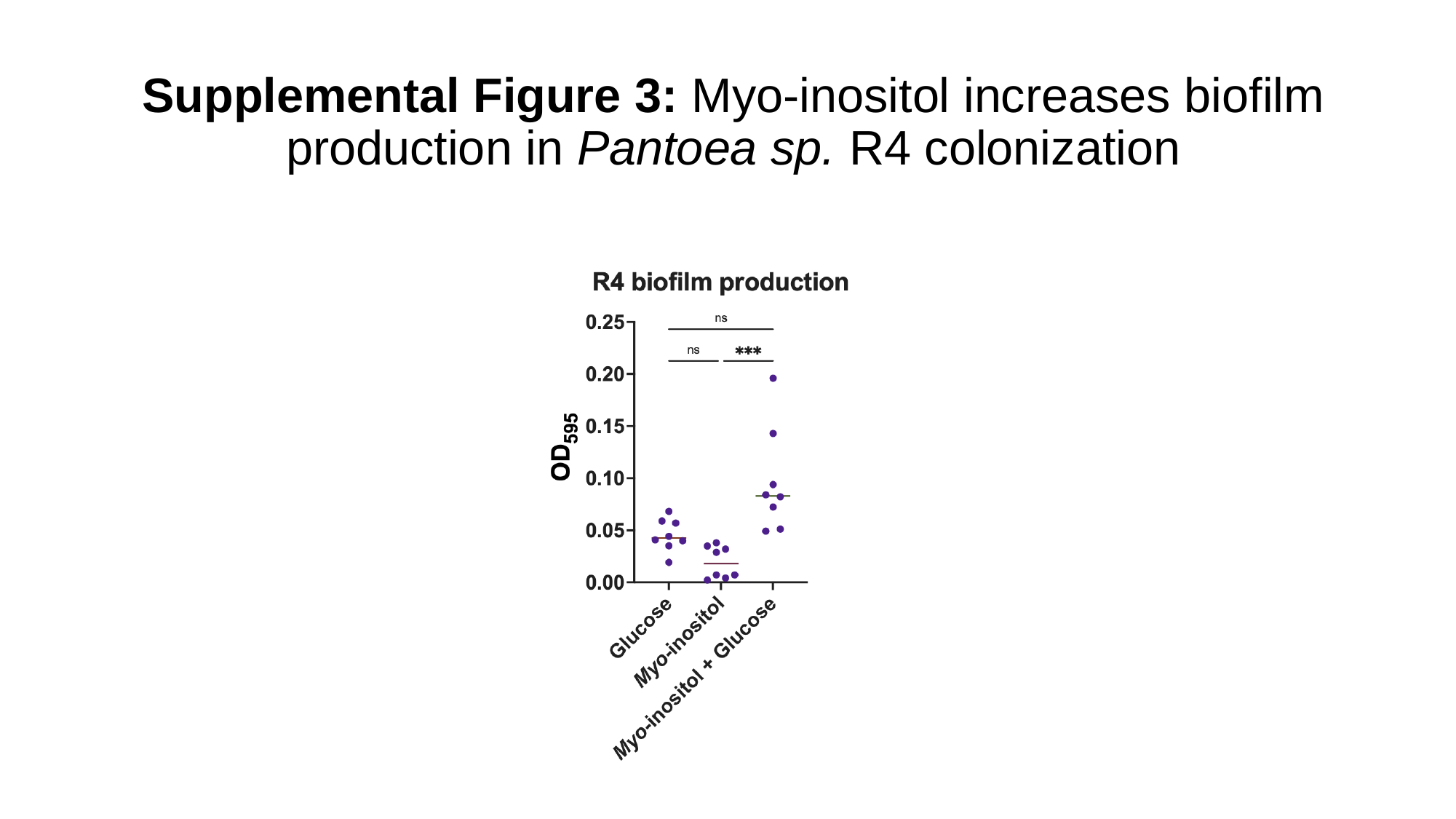

#### Supplemental Figure 3: Myo-inositol increases biofilm production in Pantoea sp. R4 colonization
